# Tuning the course of evolution on the biophysical fitness landscape of an RNA virus

**DOI:** 10.1101/090258

**Authors:** Assaf Rotem, Adrian W.R. Serohijos, Connie B. Chang, Joshua T. Wolfe, Audrey E. Fischer, Thomas S. Mehoke, Huidan Zhang, Ye Tao, W. Lloyd Ung, Jeong-Mo Choi, Abimbola O. Kolawole, Stephan A. Koehler, Susan Wu, Peter M. Thielen, Naiwen Cui, Plamen A. Demirev, Nicholas S. Giacobbi, Timothy R. Julian, Kellogg Schwab, Jeffrey S. Lin, Thomas J. Smith, James M. Pipas, Christiane E. Wobus, Andrew B. Feldman, David A. Weitz, Eugene I. Shakhnovich

**Affiliations:** School of Engineering and Applied Sciences and Department of Physics, Harvard University, 9 Oxford Street, Cambridge, MA 02138, USA; Department of Chemistry and Chemical Biology, Harvard University, 12 Oxford Street, Cambridge, MA 02138, USA; Johns Hopkins University Applied Physics Laboratory, 11100 Johns Hopkins Road, Laurel, MD 20723, USA; Department of Cell Biology, Key Laboratory of Cell Biology, Ministry of Public Health, and Key Laboratory of Medical Cell Biology, Ministry of Education, China Medical University, Shenyang 110001, China; Department of Microbiology and Immunology, University of Michigan Medical School, 1150 West Medical Center Drive, Ann Arbor, MI 48109, USA; Department of Biological Sciences, University of Pittsburgh, 4249 Fifth Avenue, Pittsburgh, PA 15260, USA.; Environmental Health Sciences and the Hopkins Water Institute, Johns Hopkins Bloomberg School of Public Health, Baltimore, Maryland 21231, USA; Department of Biochemistry and Molecular Biology, University of Texas Medical Branch at Galveston, 301 University Boulevard, Galveston, TX 77555, USA; Chemical and Biological Engineering Department, Montana State University, Bozeman, Montana, USA; Département de Biochimie et Centre Robert-Cedergren en Bioinformatique et Génomique, Université de Montréal, Quebec, Canada; Department of Emergency Medicine, Johns Hopkins Medicine, 5801 Smith Avenue, Suite 3220, Davis Building, Baltimore, MD, 21209, USA; Department of Environmental Microbiology, Swiss Federal Institute of Aquatic Science and Technology (Eawag), 8600 Dübendorf, Switzerland.

## Abstract

Predicting viral evolution remains a major challenge with profound implications for public health. Viral evolutionary pathways are determined by the fitness landscape, which maps viral genotype to fitness. However, a quantitative description of the landscape and the evolutionary forces on it remain elusive. Here, we apply a biophysical fitness model based on capsid folding stability and antibody binding affinity to predict the evolutionary pathway of norovirus escaping a neutralizing antibody. The model is validated by experimental evolution in bulk culture and in a drop-based microfluidics device, the “Evolution Chip”, which propagates millions of independent viral sub-populations. We demonstrate that along the axis of binding affinity, selection for escape variants and drift due to random mutations have the same direction. However, along folding stability, selection and drift are opposing forces whose balance is tuned by viral population size. Our results demonstrate that predictable epistatic tradeoffs shape viral evolution.

## Introduction

Responding to viral pandemics or to the emergence of new microbial pathogens is a major challenge to public health ^1^. A critical component to this response is the prediction of the course of microbial evolution. One approach to this prediction uses available genomic samples and branching patterns of genealogical trees to statistically project future dominant strains ^2^. However, this approach can only infer the likelihood that an existing viral strain will dominate, and cannot predict emergence of novel strains that might become dominant due to *de novo* beneficial mutations. It is these novel strains that are often the most virulent, posing the greatest hazard to public health. Predicting the evolution of novel strains is the key to addressing this challenge and requires knowledge of the relationship between mutations in the viral genome and the fitness for individual organisms ^3^. This relationship is the fitness landscape, which is a complex, multidimensional function; however, this can rarely be quantitatively determined. Nevertheless, it is essential for predicting selection of the most probable mutants. Moreover, the fate of mutations is also a function of population structure ^4^. In particular, population size changes the balance between the impact of random mutations on fitness and that of selection ^5^ and is thought to affect both the rate and direction of evolution ^6^. Recent studies use microbial fitness landscapes ^7-9^ or population structure ^10,11^ to predict the course of evolution, but to date, none links these elements together. Without this link, further progress in predicting the course of viral evolution is significantly hindered.

In this paper, we quantitatively determine a fitness landscape for an RNA virus subjected to the environmental pressure of a neutralizing antibody, and use it to account for the evolution of the virus under conditions that constrain population size. The experimentally measured fitness landscape is correctly described by two biophysical parameters: the thermodynamics of folding of the capsid protein and its binding to the antibody. We probe the evolution of a model norovirus both in bulk, where population size is large, and in a microfluidic evolution chip which uses small drops to perform millions of experiments ^12-16^ that probe evolution in very small population sizes. We show that the dynamics of viral adaptation is strongly dependent on population size. These results can be quantitatively described by a theoretical framework that combines protein biophysics and population genetics, providing the critical link between fitness landscape and population structure that enables prediction of evolution.

We focus in this work on Murine Norovirus (MNV), a model for human RNA viruses which are the major cause of epidemics in the world ^17-19^. MNV is a non-enveloped RNA virus that consists of 180 copies of the capsid protein assembled around a 7.5kb long positive-strand RNA genome. It mutates at ~1 base per genome per replication cycle and produces ~10^4^ progenies in a single cell infection, of which ~100 are infectious viral particles, or plaque forming units (pfu)^12^.

## Results

### Lab evolution of Norovirus in large and small population sizes

To study viral evolution, we propagate a viral isolate (MNV-1, denoted *wt*) in the presence of a neutralizing antibody (mAb6.2, ^20^) that binds to the protruding domain (P-domain) of the capsid, and prevents virus entry into the host cell ^21,22^. This set-up allows us to study the way the virus evolves to adapt to a new environment. To investigate the dynamics of this escape from the antibody, we sequence a 376 bp fragment of the genome encoding the outermost part of the P-domain (residues 281 – 412 of VP1) and follow the frequency of 37,244 unique haplotypes (**Figure 1-figure supplement 1** and **Table S1**) observed over several passages, allowing us to follow the evolution over several generations. First, we propagate *wt* in standard bulk culture conditions, using ~10^6^ virions per passage under Ab pressure (**Figure 1A**). The population is initially dominated by the *wt* (~90% of the population) with the rest of the viral quasi-species consisting of single and double mutants (**Table S1**). After 2 passages the total number of surviving viruses has decreased significantly due to the neutralizing effect of the Ab (**Figure 1-figure supplement 2**); however, three single mutants E296K (*A*), D385G (*B*), T301I (*C*) as well as their double mutants (*AB*, *AC*, *BC*) occur at higher frequencies than the other haplotypes. By the fourth passage the triple mutant *ABC*, which first arises on passage 2, occurs at a frequency even higher than the other mutant haplotypes (86%); moreover, the total number of viruses increases to levels comparable to those observed after the first passage (**Figure 1-figure supplement 2**). This suggests that *ABC* is an escape variant.

A central tenet of evolutionary theory is that the way organisms explore their fitness landscape depends on the size of their population, which controls the balance between random mutational drift (i.e., direction of *randomly arising* mutations) in the population and fitness driven selection (i.e., direction of beneficial mutations) ^4,5^. This balance determines the most likely evolutionary pathways on a given fitness landscape. Indeed, the population size may be particularly important for noroviruses where a single viral particle is sufficient to infect the host animal ^23^; thus it is possible that viruses propagate in very small populations as they adapt to a new environment prior to the emergence of an epidemic.

The consequences of a smaller population size are captured by the S. Wright's shifting balance theory ^6^, which hypothesizes that evolution can proceed more efficiently in three phases: i) a population is divided into subpopulations to weaken selection and increase drift, ii) each subpopulation evolves independently, whereupon iii) the subpopulations are mixed back into a large population and then all individuals compete. We can directly probe this hypothesis experimentally by drastically reducing population size compared to typical laboratory bulk cultures, which propagate ~10^6^ to 10^8^ viruses (**Figure 1-figure supplement 2**). To evolve viruses in small population sizes we use a novel microfluidics system, the “Evolution Chip”, which propagates ~10^6^ subpopulations of 1-10 infectious particles (pfu) in distinct and non-mixing compartments (**Figure 1B**, **Figure 1-figure supplement 3, Movie S1 and Movie S2**). The microfluidics system allows us to drastically reduce the population size without reducing the total number of viruses sampled, thereby maintaining the statistics comparable to that of a bulk experiment.

**Figure 1.**
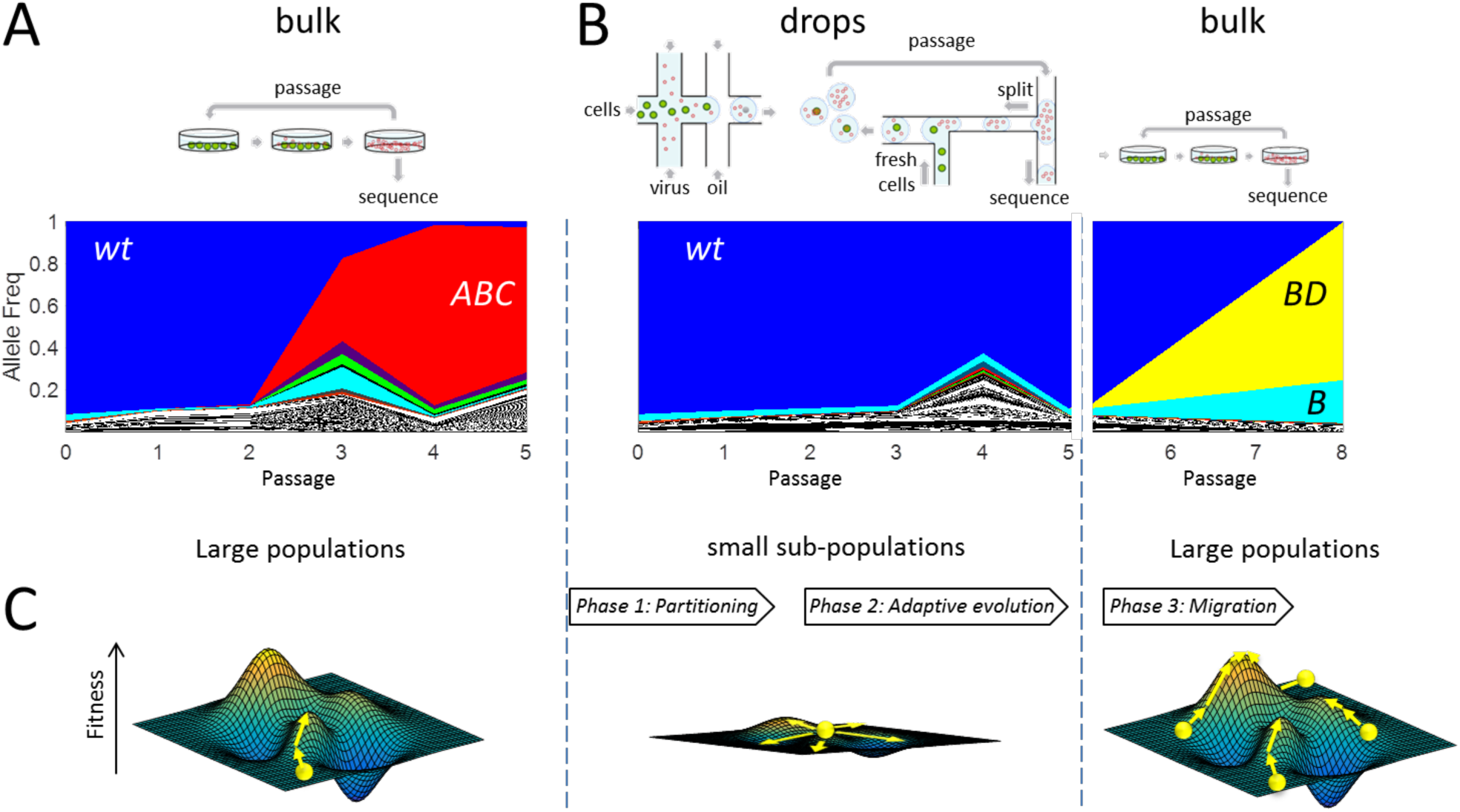
Wright's shifting balance theory (SBT)-Viral evolution in large and small population sizes. **A) Top:** 10^6^ viruses evolving against a neutralizing antibody in bulk by serial propagation. **Bottom:** The allele frequencies of 1,364 distinct P-domain haplotype sequences are plotted per passage (see also **Figure S2**). **B) Viral evolution in small populations. Top:** Experimental scheme of the three phases of Wright's SBT-*Phase 1*: 10^6^ pico-liter drops are loaded with on average 1 virus and 2 host cells per drop. *Phase 2:* viruses evolve in drops for five passages. *Phase 3:* the emulsion is broken and all lineages evolve together in bulk for 3 additional passages (frequencies for passages 6 and 7 are interpolated from passages 5 and 8, see also **Figure S3 and Movie S1, S2**). **Bottom:** The allele frequencies of 620 distinct P-domain haplotypes from B (*Phase 2, center panel, phase 3 right panel*) are plotted per passage. (**Figure S2**). **C) Exploration of the fitness landscape for the three phases of SBT**-*phase 1 (left)*: a previously adapted population (lower peak on left fitness landscape) is partitioned into smaller subpopulations. *phase 2 (middle)*: small sub-populations adapt in isolated conditions where selection pressure is reduced, allowing an extended exploration of the fitness landscape. *phase 3 (right)*: subpopulations migrate, mix and compete, evolving new and more fit variants that were explored in isolation (highest peak). Haplotype legend: A: E296K, B: D385G, C: T301I, D: A382V.

In stark contrast with the bulk experiments, amplification and hence growth of potential escape variants that sweep the population is precluded when each variant is confined in a single drop with just two host cells; as a result, the *wt* remains the dominant fraction of the observed viruses through all passages. Potential escape viruses are present, but are in complete isolation from each other, at population sizes of just a single infection event per generation. This microfluidic experiment (**Figure 1B**) implements the first two phases of the shifting balance theory ^6^: i) partitioning the population in drops to weaken selection and increase genetic drift and ii) evolving the sub-populations without competition between drops. However, the absence of competition precludes detection of potential escape variants. To overcome this, we break the emulsion after five passages in the evolution chip, mix the contents of all the drops and propagate the sample in the presence of Ab under bulk conditions for three additional passages ^26^. This enables new escape variants that evolved in isolation to take over the integrated population and facilitates their identification, isolation and characterization. This also implements the third phase of the shifting balance theory: iii) mixing the subpopulations back into a large population. Remarkably, after mixing, a double mutant, D385G-A382V (*BD*), sweeps the population; (**Figure 1B**) this is in sharp contrast with the standard bulk cultures where the triple mutant *ABC* sweeps the population.

Next, we determine if the escapees in drops are more fit than the escapees in bulk. To address this question, we engineered the mutations into the infectious clone and recovered mutant viruses. Next, we performed head-to-head competition of *wt*, *BD*, and *ABC* variants. We show in **Figure 2** the frequency of each of the three clones at the end of the competition. Indeed, *BD* and *ABC* are true escape variants since they outcompete *wt* under neutralizing antibody. However, without antibody, *wt* is more fit than both *BD* and *ABC*, which explains the observation that neither *BD* nor *ABC* spontaneously arise in serial passaging without Ab. More importantly, in the competition between *BD* and *ABC*, we find that the escapee from droplets is more fit compared to the escapee from bulk (**Figure 2**). Despite being more fit, *BD* is not detected in any of the bulk serial passaging under neutralizing Ab, while *ABC* was observed thrice. We hypothesize that the initial acquisition of the mutation D (enroute to *BD*) is limited in large populations, because epistatic interactions dictate that the virus traverse a low fitness regime, before climbing up a local fitness peak. Indeed, such unlikely pathways on the landscape can be accessed if selection is weakened by decreasing population size as Wright originally envisioned.

**Figure 2.**
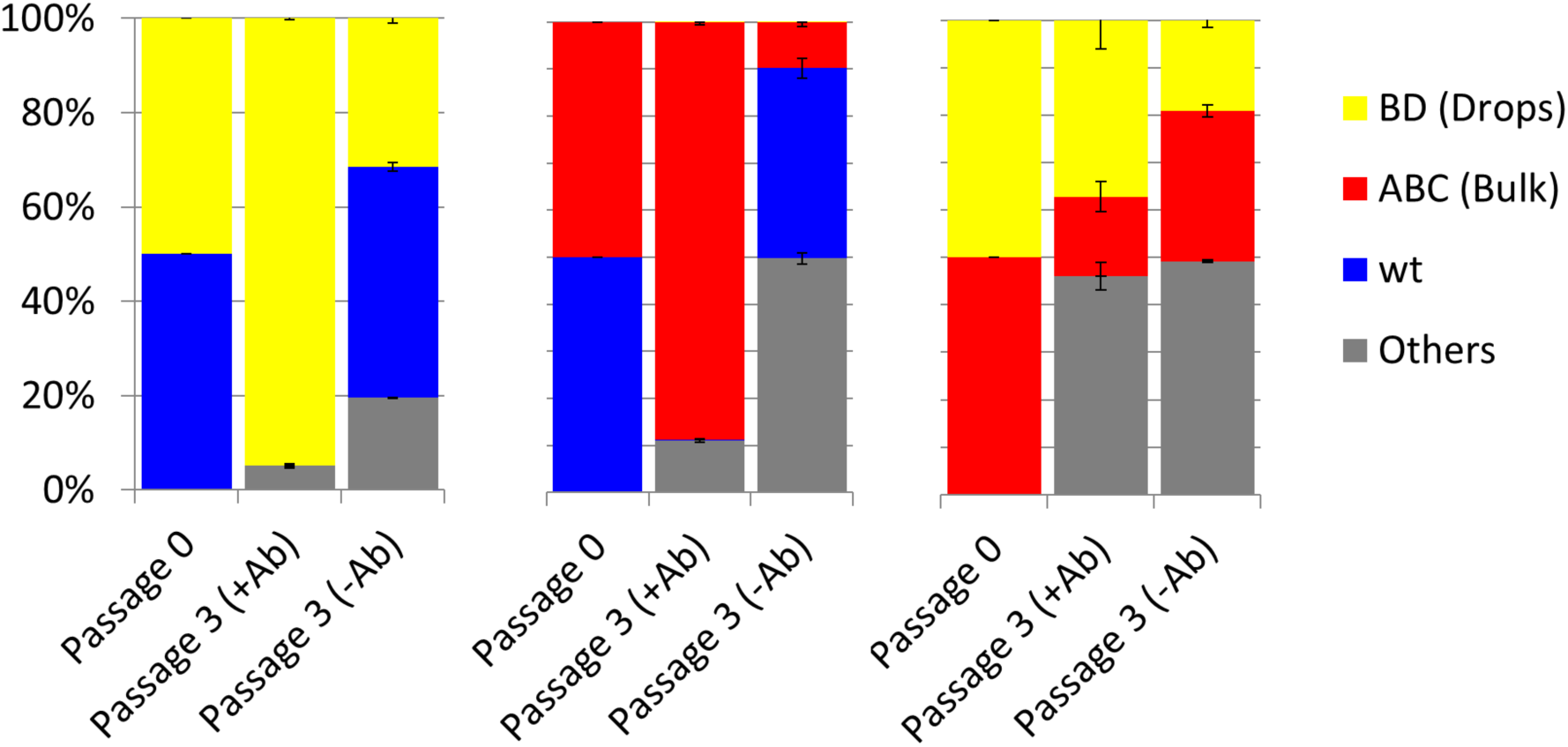
Allele frequency of the clone after 3 passages in competition assays. To perform pairwise competition of the clones, we mixed equal titers of the clone, propagate them for 3 passages, and then perform deep sequencing. Error bars are S.E. of 3 biological replicates for each measurement. See also Table S2.

### Fitness landscape of norovirus escaping an antibody is projected onto the biophysical properties of its capsid domain

These results provide strong qualitative support to Wright's shifting balance theory showing that the evolutionary dynamics and outcome dramatically depend on the population structure. To understand these dynamics quantitatively we must first develop a tractable model for the fitness landscape of the virus. In general, fitness is expected to be a complex function of multiple traits. Instead we focus on the dependence of viral fitness in the presence of a neutralizing antibody on two biophysical properties of the P-domain: The folding energy, which is a measure of stability, and the binding affinity to the antibody, which is a measure of neutralization. While the importance of binding affinity to antibody is apparent, the universal importance of protein folding stability for bacterial and viral fitness was also shown ^8,27^. This choice of variables is further supported by the fact that all the mutations of the dominant escape variants we observe in our experiments are located within the binding site between the P-domain and the Ab, as shown by mapping the mutations on the 3D structure of the *wt* P-domain in **Figure 3A**.

**Figure 3.**
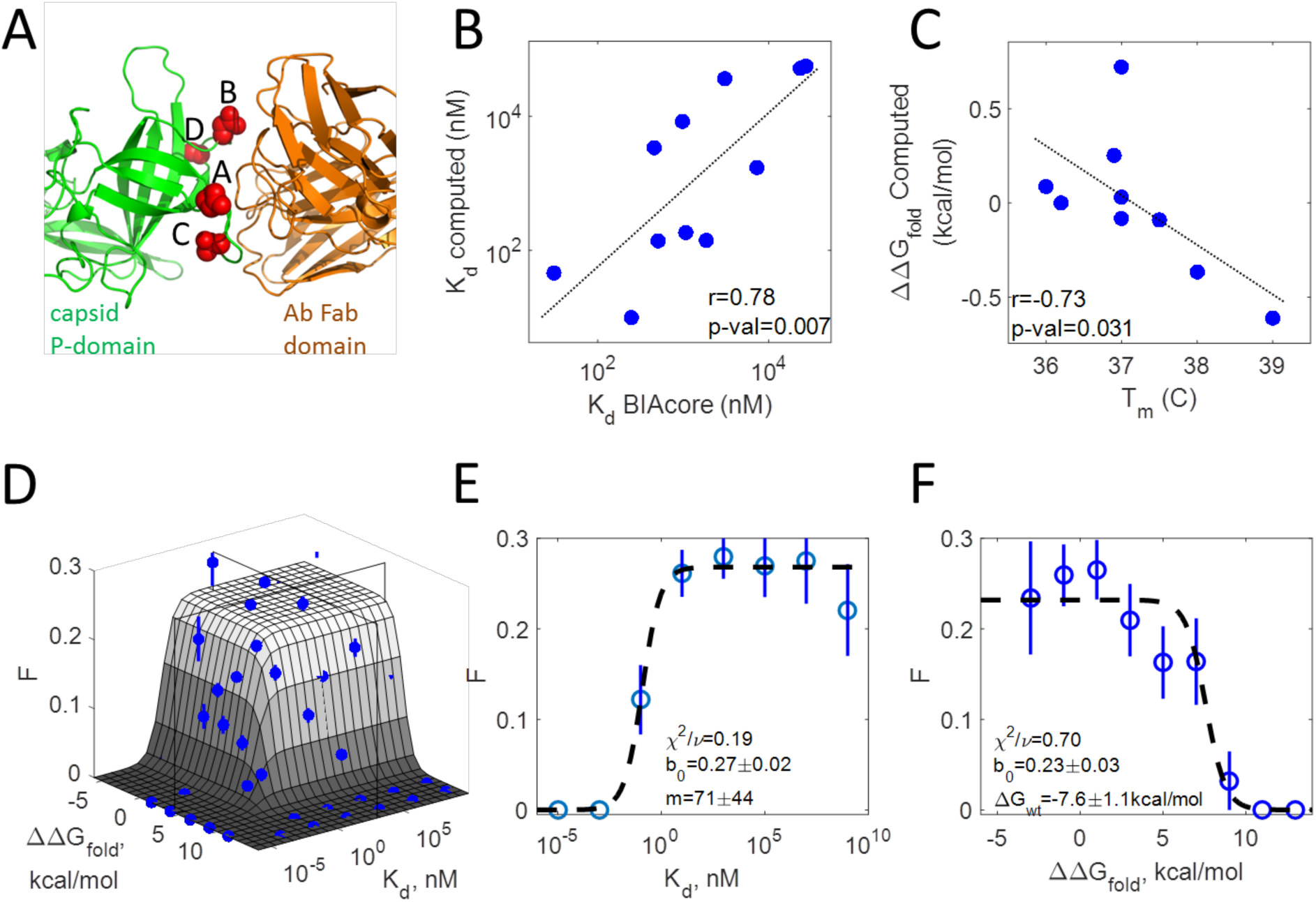
Fitness landscape of norovirus escaping a neutralizing antibody. **A)** The P-domain Antibody complex structure. The SNPs of all dominant P-domain variants (red circles) are located on the docking site of the P-domain-antibody complex (PDB ID: 3LQE). **B)** A high correlation exists between Ab dissociation constant *K_d_* that was experimentally measured using surface plasmon resonance (SPR) and the one computed from force field calculations **C)** The anti-correlation between the experimentally measured P-Domain melting temperature (*T_m_*) and the folding stability computed from force field calculations. Two outlier variants were excluded from the analysis. **D)** A 3D plot of the probability of infection *F* averaged over 2,076 distinct haplotypes binned according to their dissociation constant *K_d_* and folding stability ∆*G_fold_* (blue points) overlaid with the theoretical fit according to Eq. 1 (gray surface). Cross sections (black frames) demark the regions used for the projections in B and C. **E)** The probability of infection for all haplotypes with ∆*G_fold_*<4.5Kcal/mol (cross section parallel to *K_d_* axis in A) is projected on the *K_d_-F* plane, binned according to their *K_d_* (blue points) and overlaid with the theoretical fit to Eq. 1 (dashed line). **F)** The probability of infection for all haplotypes with *K_d_*>10^3^nM (cross section parallel to ∆*G_fold_* axis in A is projected on the ∆*G_fold_-F* plane, binned according to their ∆*G_fold_* (blue points) and overlaid with the theoretical fit to Eq. 1 (dashed line). *F* is determined from deep sequencing lysates of *in vitro* experiments in the presence of neutralizing antibody. *K_d_* and ∆*G_fold_* are estimated from mapping the haplotype mutations to the 3D structure of the capsid P-domain in complex with the neutralizing antibody.

To calculate the folding energy of the P-domain and its binding affinity to the antibody for each haplotype sequence, we use force field calculations based on the structural mapping in **Figure 3A**^26,28^ to determine the change in folding energy ∆∆*G_fold_* between the mutant and the *wt*; from this we determine ∆*G_fold_* of the mutant by adding the folding energy of the *wt*, ∆*G_wt_*. We also determine the change in binding energy, ∆∆*G_bind_*, between the mutated P-domain-Ab complex and the *wt*; from this we determine the dissociation constant *K_d_* = *K*_0_exp(*β*ΔΔ*G_bind_*) where *K_0_* is the dissociation constant of the *wt*. We test the accuracy of our calculations by comparing the calculated biophysical properties of the escape haplotypes to experimentally measured properties. To accomplish this, we express and purify the P-domain of each haplotype^29^ and measure its binding to the Ab to extract *K_d_*; we also measure the melting temperature of the P-domain, *T_m_*, which correlates inversely to ∆*G_fold_* (**Table S3** and ref. ^30^). The measured values of the biophysical properties of the dominant escape haplotypes correlate strongly with the calculated values of the same haplotypes, as shown in **Figure 3B-C**. Importantly, we reverse engineer the escape viruses with their haplotype sequences on the background of the *wt* for the rest of the virus and confirm that the observed mutations in the P-domain are directly responsible for their increase in fitness both *in vitro* (**Figure 4A** and **Figure 4-figure supplement 1A**) and *in vivo* in mice (**Figure 4B** and **Figure 4-figure supplement 1B**); thus, our biophysical variables are relevant for viral fitness inside the real animal host.

**Figure 4.**
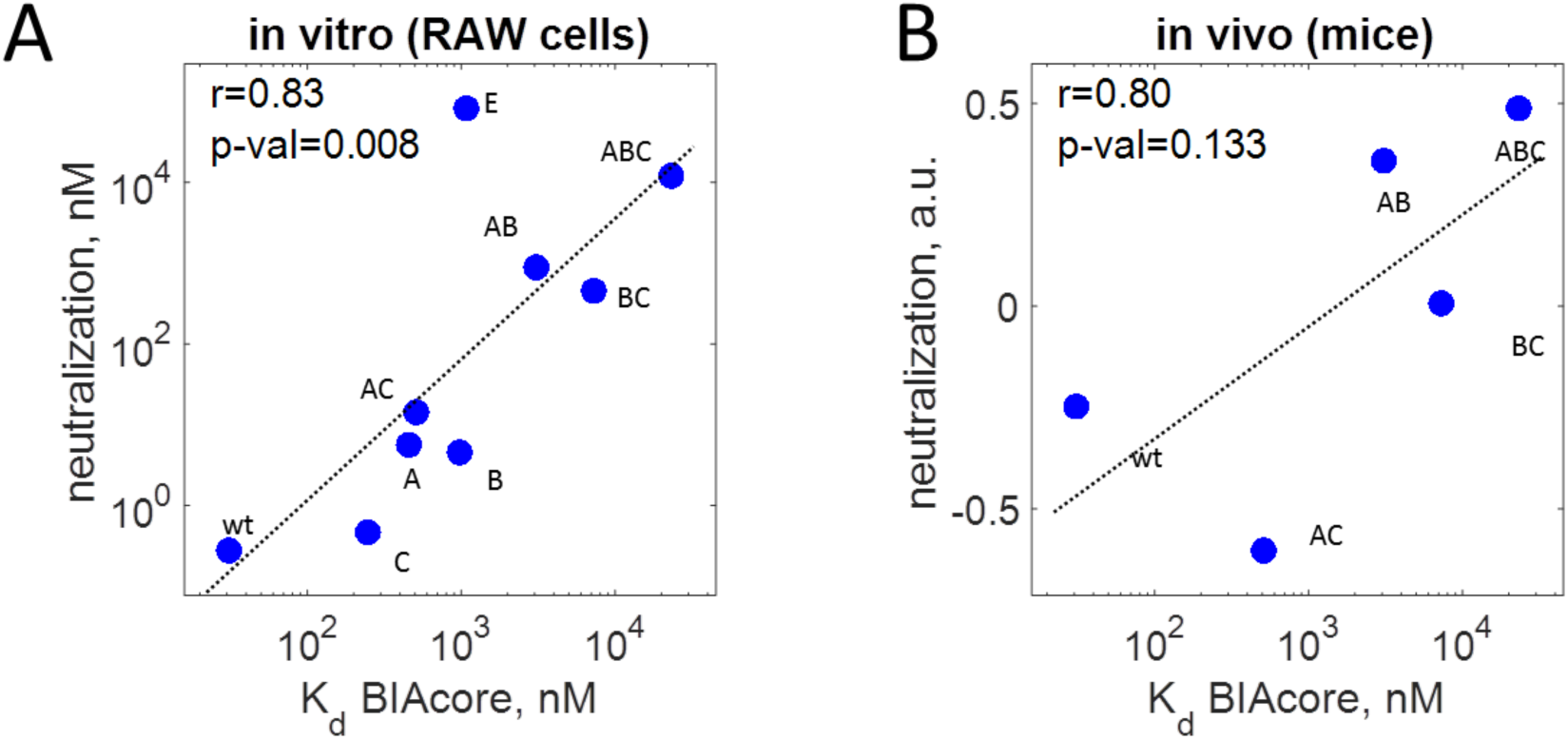
MNV-1 neutralization versus binding affinity of the P-domain to neutralizing antibody. **A)** *in vitro* neutralization of dominant haplotypes correlates to their *K_d_* and the average ratio between them is ~120, in good agreement with the modeled value of *m*≈70. E: L386F **B)** *in vivo* neutralization of dominant haplotypes in mice correlates to their *K_d_*.

The biophysical fitness landscape describes the dependence of viral fitness in the presence of a neutralizing antibody on ∆*G_fold_* and 1/(*mKd*), where the parameter *m* accounts for the multiple binding sites of the capsid. To formulate the viral fitness we assume that the *wt* P-domain occurs in three specific states: folded and unbound, folded and bound, and unfolded (which is always unbound). The virus infects only when the P-domain is folded and unbound, hence, we can express the viral infectivity *F* at a given concentration of antibody [*Ab*] as^31^:

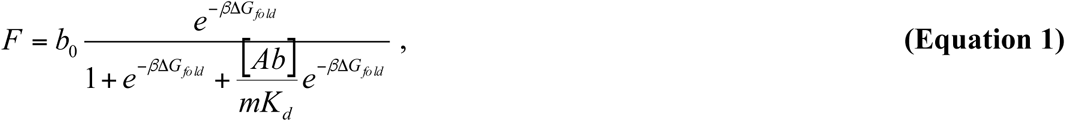

where the numerator is the Boltzmann probability of being folded and unbound and the denominator is the partition function that sums over the probability of all three states, and *β*=1/*kBT* where *k_B_* is the Boltzmann constant and *T* is the temperature. The function *F* has two regimes as shown by the surface in **Figure 3D**. For low binding affinities and stable P-domain structures, viruses are expected to infect host cells at some fixed probability, *0<b_0_<1*, determined by the average effect of all remaining viral properties on the infection process, and *F*=*b_0_.* By contrast, when the binding to the Ab is strong or when the P-domain is unstable, the virus cannot infect its host and *F=0*.

To compare the model to experiment, we use sequencing data to determine the growth rate of each virus from the change in genome haplotype frequencies between successive generations ^9^. The growth rates distribute into two distinct groups with 87% of haplotypes exhibiting little or no growth and the rest exhibiting considerably larger growth. We take the first group to be non-infective, and take the second group to be infective (**Figure 3-figure supplement 1A**). For each haplotype sequence, we map the mutations to the 3D structure of the *wt* P-domain and use Eris force field calculations to determine the change in folding energy ΔΔ*G_fold_* between the mutant and the *wt;* from this we determine stability of the mutant Δ*G_fold_*=Δ*G_fold_*,*wt*+ΔΔ*G_fold_*, where Δ*G_wt_* is the folding energy of the *wt*. We also determine the change in binding energy, ΔΔ*G_bind_*, between the mutated P-domain-Ab complex and the *wt;* from this we determine the dissociation constant *K_d_=K_0_e^βΔΔG^bind* where *K_0_* is the dissociation constant of the *wt*. We bin the haplotypes using ΔΔ*G_fold_* and *K_d_* and calculate *F* for each bin from the fraction of infective haplotypes. This binning exploits the large number of unique haplotypes to reduce the effects of errors in the calculations and of contributions from other biophysical properties. We fit the model by varying the three unknown parameters, *b_0_*, ∆*G_wt_* and the multiplier *m*.

We obtain excellent agreement between the model and the data, as shown by the dashed line in **Figure 3E-F**. The infectivity of haplotypes is zero at low *K_d_* or high ∆*G_fold_*, while at high *K_d_* and low ∆*G_fold_*, the landscape plateaus at *F*~0.25 independent of either of the biophysical coordinates. The value of 1/*m*≈1.4% obtained from the fit reflects the fact that only about 3 of the 180 P-domains on the capsid have to be blocked by the Ab to prevent infection (**Figure 3E** and ref. ^12^). The value of *F ~0.25* at the plateau is significantly less than the expected value of *b_0_*=1; this points to the role of factors not included in the model, such as the interaction of the capsid with the host-cell receptor, in successful infection. In the absence of the neutralizing antibody, we do not expect the fitness landscape to depend on *K_d_*, which is indeed the case (**Figure 3-figure supplement 2**). Altogether, these results demonstrate that binding and folding energies are very good predictors of viral extinction; however they are less successful in predicting infectivity. This suggests that antibody escape and folding stability of the capsid protein are necessary but not sufficient for viral infection.

### Most likely pathways on the fitness landscape predicted by protein biophysics and population genetics

To determine how the virus evolves on the fitness landscape, we use population genetics theory and calculate the ratio dN/dS, where *dN* is the rate of non-synonymous evolutionary rate and *dS* is the synonymous evolutionary rate ^32^ (For greater details, see section on Balance between selection and mutational drift on the fitness landscape in the Materials and Methods). As a rule, mutants affect the population more in regions where non-synonymous mutations are beneficial (high dN/dS), driving it down the gradient of dN/dS towards regions where non-synonymous mutations are deleterious (low dN/dS) and thus do not affect the population. The dN/dS ratio is a function of population size and of the fitness effect of a mutation or the selection coefficient *s=(F_mutant_-F_widltype_)/F_widltype_* ^32,33^. Using the fitness landscape from Equation 1, we calculate the dN/dS ratio on the viral fitness landscape and how it depends on population size (see Methods and ^34,35^ for details). In large population sizes, dN/dS exhibits a strong gradient towards high *K_d_* and high folding stability (**Figure 5A**). Consequently, a *wt* population that is initially unstable and neutralized by Ab is expected to evolve resistance by increasing both *K_d_* and stability. However, in small population sizes the gradient of dN/dS is directed only towards high *K_d_* and the same initially unstable population will evolve resistance to Ab without increasing folding stability (**Figure 5B**).

The effect of population size on the course of evolution can be explained by analyzing the balance between selection and mutational drift, the two forces driving evolution. The direction of drift and selection along the trait of folding stability and binding affinity to the antibody can be inferred from protein engineering and systematic studies on effects of random mutations on proteins. Along the axis of folding stability Δ*G_fold_*, beneficial mutations increase folding stability, but random de novo mutations in proteins tend to decrease stability^36-39^. Thus, selection and drift act in opposite directions (**Figure 5A-B**, arrows), leading to mutation-selection balance. Along the axis of binding affinity *K_d_*, beneficial mutations for the virus lead to escape from Ab (towards high *K_d_*), and random mutations on protein interfaces perturb binding (also towards high *K_d_*). Thus, selection and drift act along the same direction (**Figure 5A-B**, arrows).

To validate the difference in expected pathways to antibody resistance based on population size, we simulate the trajectory for 4 sequential fixations of a single mutation starting from the position of the *wt* virus population on the viral fitness landscape (see ^31^ for details). For large *N*, the simulations show that the norovirus population increases both its folding stability and dissociation constant, *K_d_*, as it escapes the neutralizing antibody, following the trajectory shown by the solid points in **Figure 5A-B** (see also **Figure 5-figure supplement 1** for simulations in the polyclonal regime). After 4 sequential single mutations the model shows a rise of 5 orders of magnitude in *K_d_* and an increase of 3kcal/mol in ∆*G_fold_* towards stabilization corresponding to a change of 9°C in *T_m_*. On the other hand, for small *N* the simulations show that the norovirus population increases its *K_d_* to escape the antibody but exhibits only a small change in folding stability of less than 1kcal/mol, corresponding to an increase in *T_m_* of about 2°C, as shown by the trajectory in **Figure 5B** (see also **Figure 5-figure supplement 1**).

**Figure 5.**
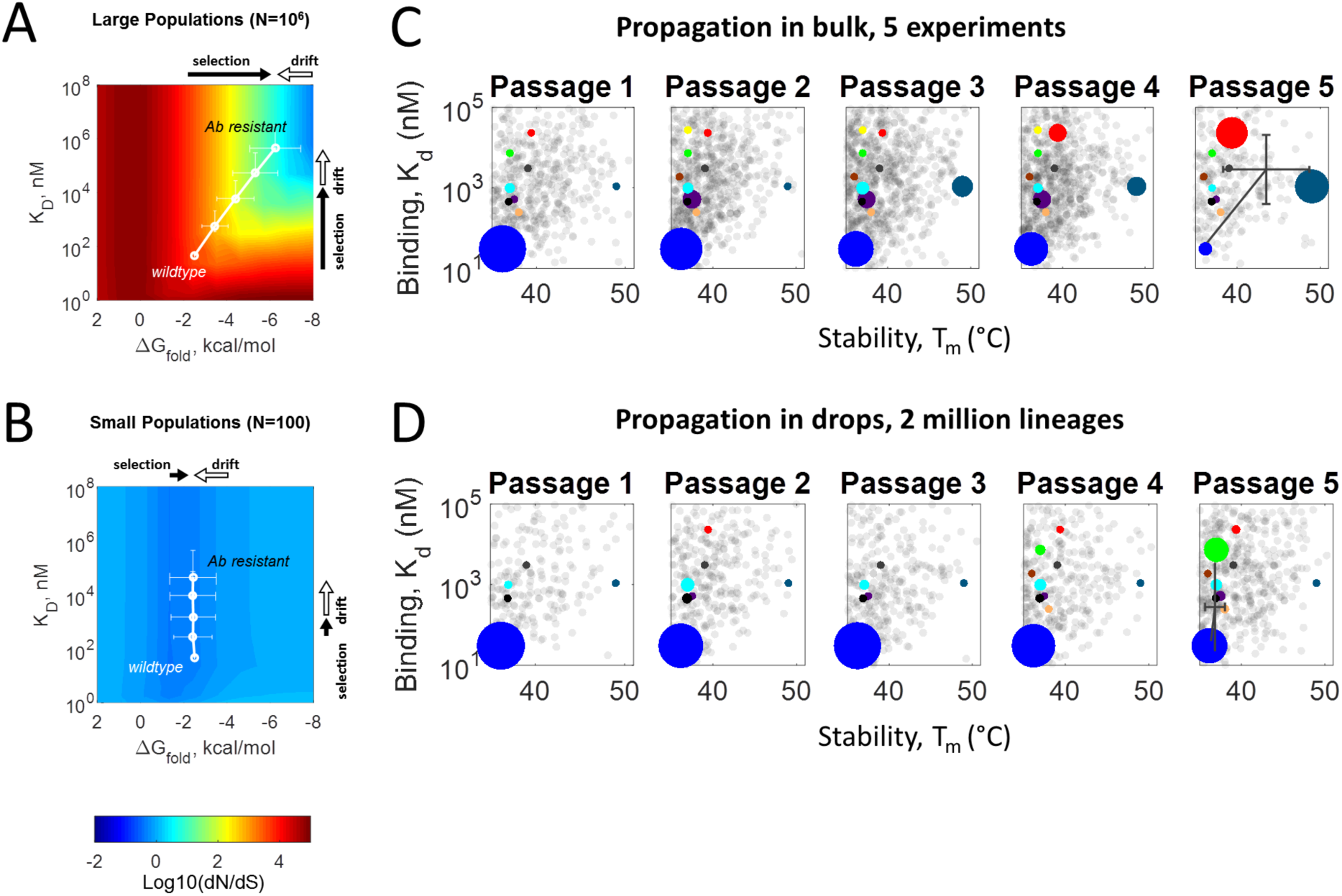
Most likely pathways on the fitness landscape predicted by protein biophysics and population genetics. **A,B)** Average stringency of selection for several population sizes (see Text and). For large population sizes, the increase in *K_d_* is strongly coupled to the increase in *T_m_*. However, for small population sizes, the selection for *K_d_* is decoupled from the selection for folding stability. The white lines are the predicted trajectories from forward evolutionary simulations of an MNV population escaping an Ab, but with a P-domain which is unstable. Each trajectory is the average of 1000 independent simulations. The direction of selection (black arrows) is towards greater folding stability and weaker affinity to the antibody. Selection is strong when the P-domain is unstable and/or is tightly bound to the Ab. Selection pressure is approximately zero when the fitness landscape is flat (neutral). Along the direction of folding stability, most random mutations are destabilizing which lead to a mutational drift (white arrows) towards protein destabilization. Along binding affinity axis, most random mutations perturb the protein-protein interaction that leads to a mutational drift towards weaker binding. **C,D)** Density plots of all haplotypes grouped according to passages. Color circles denote dominant haplotypes whose biophysical properties were measured, while the remaining gray circles denote haplotypes whose biophysical properties were calculated. The size of the circle represents the allele frequency of each haplotype. Gray cross at passage 5 denotes mean and s.d. of all haplotypes.

To compare the theory to experiment we plot the position of viral haplotypes evolving in 5 independent bulk passaging experiments as a function of *K_d_* and *T_m_*, denoting their frequencies by the size of the circles for each passage. To that end we include the calculated biophysical values *K_d_* and *T_m_* (gray haplotypes), using the correlation curve in **Figure 3B** to relate between ∆*G_fold_* and *T_m_*, as well as the biophysically measured values for select haplotypes (colored symbols). There is a clear trajectory as the intermediate variants evolve, having increasingly weaker affinities and higher *T_m_*, with the escape variants at passage 5 ultimately having the weakest affinity, with an overall average of *K_d_* ~3,000 nM and the highest P-domain stability with an average of *T_m_*~43.5°C, as shown in **Figure 5C**. We also plot the position of viral haplotypes evolving in ~2 million independent drop passaging experiments. We observe a clear trajectory of increasingly weaker affinities while maintaining original *T_m_*, with the escape variants at passage 5 ultimately having an overall average of *K_d_*~300 nM and *T_m_*~36.8°C, as shown in **Figure 5D**. The direction of these trajectories parallels that of the simulated trajectories, supporting our choice of these explicit biophysical properties as valid and useful coordinates for the fitness landscape. This also provides us a theoretical framework with which to interpret new experimental data and to test fundamental concepts of evolutionary theory.

## Discussion

It is a central concept in evolutionary biology theory that the population size determines the balance between two evolutionary forces, selection and mutational drift ^5,34^. The results presented here provide direct experimental evidence in support of this central concept. Moreover, the role of Wright's shifting balance theory in real evolution has been contentious and controversial because of lack of direct experimental demonstration^40^. Using the segregation in the Evolution Chip, we show that isolated viral populations starting at a fitness valley in the presence of a neutralizing Ab are able to explore unlikely pathways on the adaptive fitness landscape and, following migration, shift the whole population to new fitness peaks (**Figure 1C**); this is a direct demonstration of the shifting balance theory.

The ability to change the course of evolution using the Evolution Chip and to predict its direction on the biophysical fitness landscape with the parameters of protein stability and binding may be helpful in addressing pandemics. These biophysical parameters are more generally applicable to other viruses; for example, both binding ^41^ and folding stability ^8^ are relevant traits for the evolution of an influenza virus in the presence of a neutralizing antibody. Prediction of future diversification of circulating viral mutants is key to developing and potentially fielding an effective vaccine prior to, or in the early stages of a pandemic. The methodology presented here may assist in proactive exploration of viral diversification to inform selection of novel viruses prior to their natural emergence, for developing viral therapeutics.

## Materials and Methods

### Microfluidic devices

We fabricate polydimethylsiloxane (PDMS) devices using photolithography and coat them with fluorophilic Aquapel (Rider, MA, USA) to prevent wetting of drops on the channel walls. Electrodes are fabricated on chip using low melting temperature solder ^42^. The designs used to fabricate the devices are available in ACAD format (**Supplementary File S1**). We use OEM syringe pumps (KD Scientific, MA, USA) to drive the fluidics and a fast camera (HiSpec1, Fastec Imaging,USA) to image encapsulation and drop fusion.

### Reagents

For the inert carrier oil we use HFE-7500 (3M, USA) with 1% w/w of a block co-polymer surfactant of perfluorinated polyethers (PFPE) and polyethyleneglycol (PEG)^43^. A compatible surfactant is available commercially (008-FluoroSurfactant, Ran Biotechnologies, USA). To separate the emulsion, we use a commercially available demulsifier (1H,1H,2H,2H-perfluoro-1-octanol, CAS # 647-42-7). Other chemicals were purchased from Sigma Aldrich (St. Louis, MO).

### Clones and Antibodies

The plaque-purified MNV-1 clone (GV/MNV-1/2002/USA) MNV-1.CW3 ^44^ (referred herein as *wt*) was used at passage 6 for in vitro passaging experiments. Recombinant MNV-1 viruses containing P-domain point mutants E296K (*A*), D385G (*B*), T301I (*C*), A382V (*D*), L386F (*E*), E296K-T301I (*AC*), E296K-D385G (*AB*), T301I-D385G (*BC*), A382V-D385G (*BD*) and E296K-T301I-D385G (*ABC*) were generated as previously described ^20^. The isotype control IgG directed against Coxsackievirus B4 (CV) (clone 204-4) was purchased from ATCC (HB 185). Anti-MNV-1 monoclonal antibody (mAb) A6.2 (IgG2a) was grown in Bioreactor CELLine CL 1000 flasks (Sigma-Aldrich) following the manufacturer's recommendations. Antibodies were purified over a HiTrap protein A column (GE Healthcare) according to the manufacturer's instruction, dialyzed against PBS, and stored at-20 °C.

### MNV-1 P-domain mutant expression and purification

All recombinant proteins was expressed and purified as previously reported ^21^ with some modifications. Briefly, the P domain of MNV-1 (residues 225 to 541) was cloned into a pUC57 expression vector with NH2-terminal 6-histidine tag. The protein was expressed overnight at 20°C in Escherichia coli. The cells were subsequently lysed, and the protein was obtained from the supernatant, two steps purification by Ni column and by gel filtration on a Superdex 75 column (GE Healthcare). The proteins were dialyzed overnight at 4°C against a phosphate buffer (pH=7.8 and 20 mM NaCl).

### Folding stability measurement by thermofluorescence

Thermal denaturation was carried out using melt-curve module of BioRad CFX96, and Sypro Orange dye as a probe for unfolding as described earlier ^45^. The dye was added to the final concentration of 5× in a 25 *µ*l reaction volume containing 4 *µ*M of protein in 10 mM sodium phosphate buffer pH 7.8. To control for concentration dependence, we also performed the experiment using 2 *µ*M concentration of protein. The *T_m_* is estimated as the extremum of the derivative of the fluorescence signal. We report the average *T_m_* of eight technical repeats, four for 4 *µ*M of protein and another four for 2 *µ*M of protein.

### Binding kinetics by surface plasmon resonance

Realtime biomolecular analysis was performed by surface plasmon resonance (SPR) using a BIAcore 3000 instrument equipped with nitrilotriacetic acid (NTA) sensor chip. Purified monoclonal antibody A6.2 was immobilized on the surface while the MNV-1 P-domain variants were the analytes. To perform single kinetic measurement, we (1) Quickinjected 20 *µ*l of buffer (10 mM sodium phosphate buffer pH 7.8 with 0.005% surfactant to prevent minimizing nonspecific interaction); (2) Kinjected 50 uL of protein with 100 s dissociation; (3) Quickinjected 10 *µ*l of regeneration buffer (pH 2.0); and (4) Quickinjected 20 *µ*l of phosphate buffer. For each MNV variant, we performed kinetic measurements with concentrations in the range of 0.5 to 100 *µ*M. Data analysis was conducted with BIAevaluation package. Curve fittings were done with the 1:1 Langmuir binding model.

### Cell Culture

RAW 264.7 (murine macrophage) cells were purchased from ATCC (Manassas, VA) and maintained as described previously ^21,46^. Adherent cell culture medium (RAW medium) contains Dulbecco's Modified Eagle's Medium, 4 mM L-glutamine, 100 *µ*g/mL penicillin, 100 *µ*g/mL streptomycin, 10 mM HEPES, 10% heat-inactivated fetal bovine serum. The RAW 264.7 were adapted to suspension culture in spinner flasks for these experiments, for compatibility with drop-based microfluidics. Suspension cell culture medium (Suspension Medium) contains adherent RAW medium supplemented with sodium bicarbonate (7.5%). BSRT7 (BHK cells expressing T7 polymerase) cells were cultured in DMEM supplemented with 10% heat-inactivated fetal bovine serum, 2 mM L-glutamine, 100 U/ml penicillin and 100 *µ*g/ml streptomycin^47^. For all viral infection experiments, suspension RAW 264.7 cells were suspended in Enhanced Suspension Medium (ESM) comprised of Suspension Medium supplemented with 15% optiprep.

### Viral evolution in bulk

RAW 264.7 suspension cells are centrifuged for 5 min at 3,000 rpm and re-suspended in fresh medium at a concentration of 6×10^6^ cells/mL. In antibody neutralization experiments, virus is first incubated for 30 min at 37°C with mAb A6.2 in 200 *µ*L PBS prior to dilution into ESM at 4×10^6^ pfu/mL, such that the final Ab concentration is 8.57 nM. One mL of cell suspension and 1 mL of virus suspension are mixed in a single well of a 12-well dish containing a sterile stir bar and incubated on stir plate in a 37°C incubator, 5% CO2, for 24 hrs. To passage viral progeny to the next generation (the progeny of the first inoculum is considered passage 0 (P0)), 250 uL of the supernatant from cell lysates of the previous generation is supplemented to 2 mL of a fresh suspension of 2×10^6^ RAW cells/mL in ESM in a 12-well dish containing a sterile stir bar and incubated on the stir plate in a 37°C incubator, 5% CO2, for 24 hrs. In antibody neutralization experiments, 8.57 nM mAb A6.2 is supplemented to the fresh suspension of RAW cells prior to passaging. Cell lysates were harvested by 2 rounds of freeze/thaw and centrifugation for 5 min at 5,000 × g.

### Viral evolution in drops

To evolve viruses in small population sizes we use a novel microfluidics system, the “Evolution Chip”, which propagates ~10^6^ subpopulations of 1-10 infectious particles (pfu) in distinct and non-mixing compartments. We encapsulate ~2 host cells and ~1 pfu in 100*µ*m diameter aqueous drops in inert oil at a rate of millions per hour. The resultant emulsion is incubated under physiological conditions for one viral life cycle (~24 hours). Successful replication of the virus leads to the death of the host cell and the release of viral progeny within the drop. Thus, to enable continued passage, each drop is split and ~10% of its volume, containing viral progeny from the previous generation, is merged with a new drop containing a fresh host cell for the next generation.

Suspension-adapted RAW 264.7 cells were centrifuged for 5 min at 3,000 rpm, re-suspended in ESM at 8×10^6^ cells/mL. The suspension of cells is co-flowed at a 1:1 ratio with MNV-1 virus diluted into ESM at 4×10^6^ pfu/mL. The two aqueous phases-cell suspension and buffer-meet immediately before passing through the microfluidic drop making junction so that they only mix inside the 100um drops containing them (**Supplementary Movie 1**). For the continuous phase dispersing the drops, we use HFE-7500 Oil with 1% surfactant. Typical flow rates are 8 mL/hr for the oil and 2 mL/hr for cells and virus. Drops were collected in 1.5 mL tubes and incubated at 37°C, in 5% CO_2_. Following 18-24 hr incubation at 37°C, drops are re-injected into the “evolution-chip” microfluidic device where ~10% of their volume is split off and fused with freshly formed drops containing freshly prepared suspension of RAW cells in ESM. The synchronization in the device ensures that in >95% of cases, the split from one drop fuses with exactly one newly formed drop, enabling the viral lineages to propagate in isolation. Typical flow rates are 4 mL/hr for the oil, 1 mL/hr for the re-injected drops, and 1 mL/hr for the fresh cells. The newly formed drops are collected in 1.5 mL tubes and incubated at 37°C, in 5% CO_2_, while the content of the old, split drops are extracted for analysis (The progeny of the first encapsulation is considered passage 0 (P0)). In antibody neutralization experiments, 8.57 nM mAb A6.2 is supplemented to the fresh suspension of RAW cells prior to re-injection of drops.

### Head to head competitions between viral strains

Next, we determine if the escapees in drops are more fit than the escapees in bulk. To address this question, we performed head-to-head competition of *wt*, *BD*, and *ABC* strains. We mixed equal viral titers of each pair of strains, passaged them for 3 rounds and then performed deep sequencing. The competition was performed under two conditions, with and without neutralizing antibody, and over three replicates. The sequencing results of the competitions are given in **Table S2**.

RAW 264.7 suspension cells are centrifuged for 5 min at 3,000 rpm and 2 mL of a fresh suspensions of 1×10^6^ RAW cells/mL in RAW medium is dispensed in 6-well plates. In antibody neutralization experiments, virus is first incubated for 30 min at 37°C with mAb A6.2 in 200 *µ*L PBS prior to adding it into the wells, such that the final Ab concentration is 0.08 nM. Cells and virus are incubated in a 37°C incubator, 5% CO_2_, for 48 hrs. To passage viral progeny to the next generation (the progeny of the first inoculum is considered passage 0 (P0)), 200 uL of the supernatant from cell lysates of the previous generation is supplemented to 2 mL of a fresh suspension of 1×10^6^ RAW cells/mL in RAW medium in a 6-well dish and incubated in a 37°C incubator, 5% CO_2_, for 24 hrs. In antibody neutralization experiments, 0.08 nM mAb A6.2 is supplemented to the fresh suspension of RAW cells prior to passaging. Cell lysates were harvested by 2 rounds of freeze/thaw and centrifugation for 5 min at 5000 × g.

### In-vitro neutralization measurements

To measure neutralization in-vitro we followed Fischer et. al ^12^ to obtain a neutralization curve which we then fit by the equation 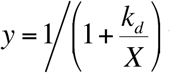 where *y* is the viral titer, *X* is the *Ab* concentration and *k_d_* is the fitting parameter.

### In-vivo neutralization measurements

Mouse studies were performed in accordance with local and federal guidelines as outlined in the “Guide for the Care and Use of Laboratory Animals” of the National Institutes of Health. Protocols were approved by the University of Michigan Committee on Use and Care of Animals (UCUCA Number: 09710). Viral strains were neutralized in STAT^-/-^ mice injected with 500ug mAb A6.2 and compared to their infection in mice injected with an isotype as described in ^48^. The decrease in viral titers was first standardized for each tissue across viral strains, before the average over all tissues within each strain was taken as its final neutralization score. The study was performed in biological triplicates. No randomization or blinding was used.

### Measurement of infectious virus titer

Viral titer was determined by either plaque assay, as described previously ^49^, or by TCID50 assay, as described previously ^50^. TCID 50% infectivity was translated to plaque forming unit (pfu) using the conversion of 0.7 TCID50 units to 1 pfu.

### Quantification of viral genome

Viral RNA isolated from experimental samples by MagMAX Viral RNA Isolation Kit (Life Technologies) was evaluated by qPCR using MNV-1-specific primer/probe sequences (Forward: GTGCGCAACACAGAGAAACG, Reverse: CGGGCTGAGCTTCCTGC, and probe: FAM/CTAGTGTCTCCTTTGGAGCACCTA/TAMARA) ^21^. Takara One Step reagent (Mountain View, CA) was used to measure the genome copy of viral samples. qRT-PCR was performed on ABI real time PCR machine (OneStep Plus) using the following thermal cycling parameters: 5 min at 42° C and 10 sec at 95° C, 40 cycles of 5 sec at 95° C and 34 sec at 60° C. Titered MNV-1 viral stock and PicoGreen (quBit)-quantitated pT7 MNV 3'RZ plasmid ^50^ were used as standards for pfu/mL and genome copies/mL analysis, respectively.

### Amplicon preparation and sequencing

Viral RNA was isolated using the MagMax viral RNA kit (Life Technologies), then subjected to DNAse digest using DNAfree Turbo DNAse (Life Technologies). SuperScript III reverse transcriptase (Life Technologies) was then used to convert vRNA to first strand DNA with specific primers tiling the entirety of MNV-1 at ~1kb increments using the following primers:

AGCCGATCACAGGCTCCTTGGC
CCATGTTGGATAAGAGGGCTGGC
ACGCACTTCCTCAACTCAGCCG
GGCCATGCTGATCCTGGCCA
CCACCAGGATGCCATCCGAGA
GTCGACATCAGCGCGTGGTATGA
CAACAGGGTGGGCACCACGTC
CAACAACAGGGCTCTCAGCATAAACCAG

Library preparation for Illumina sequencing on the miSeq platform was carried out in accordance with Illumina tech note 15044223 Rev. A using KAPA HotStart HiFi DNA polymerase 2× master mix (KAPA Biotechnologies), with primers for MNV-1 capsid amplification corresponding to MNV-1 VP1 nucleotides 848-1275 (Forward: TCGTCGGCAGCGTCAGATGTGTATAAGAGACAG**GTTCATGGGTGTCCTGCTTT**, Reverse: GTCTCGTGGGCTCGGAGATGTGTATAAGAGACAG**GGGGAGAAAGGGACCAATT**; gene specific portion in bold). After purification with Ampure XP (Beckman Genomics), samples were quantified using Qubit high sensitivity reagents (Life Technologies), then pooled in equimolar ratios assuming specific amplification during adapter addition. Final pools of up to 96 libraries were quantified using high sensitivity Bioanalyzer reagents (Agilent) prior to sequencing 2X250 bp long paired end reads of the amplicons with miSeq (v2 chemistry, miSeq control software v2.3.0.3). Samples were supplemented with 10% phiX control library (Illumina). All reagents were used according to manufacturer's recommendations, and obtained from Life Technologies unless otherwise noted.

### Amplicon sequencing analysis

Paired end reads were obtained from the MiSeq after going through the MiSeq Reporter Generate FASTQ Workflow, which demultiplexes the raw data by MID pair (removing adapter sequences in the process), and creates a separate pair of output files for each paired-end sample. These paired-end reads are aligned to the MNV-1 reference sequence (RefSeq: NC_008311) using bowtie2 (http://bowtie-bio.sourceforge.net/bowtie2/) and only reads that include the amplicon range (nucleotides 5918 to 6293 of MNV-1) without insertions and deletions are analyzed. SNPs in each of the remaining reads are saved for further analysis together with their Illumina reported quality. SNPs in overlapping regions of the paired end reads are called using the read with the higher quality. If the call in both forward and reverse reads match, the quality of SNP is taken as the sum of qualities, otherwise, the difference between the higher and lower quality is taken. Sequences with more than 20 substitutions were discarded and the remaining reads were then clustered into unique candidate haplotypes and tested for statistical significance using a hypothesis test. The null hypothesis presumed that a haplotype candidate is generated by sequencing errors in the reading of one of its “potential ancestors”. Potential ancestors are haplotypes that are both “nearest neighbors”-have the smallest hamming distance to the candidate-and are more frequent-have more reads than the candidate. We used a simple generative model using a base-calling substitution rate equal to the lowest read quality recorded for any of the relevant substitutions in the candidate reads and then calculated the probability that the observed candidate was derived from this distribution (see next section on Determination of Significance).

### Determination of significance from sequencing

Initially, the most abundant haplotype is assigned P-value=0. Haplotypes are tested in descending order of their frequency. For each haplotype, a group of “potential ancestor haplotypes” was defined as those haplotypes that are both “nearest neighbors”-have the smallest hamming distance to it-and are more frequent-have more reads than this haplotype. The single event probability that a copy of the candidate haplotype was generated while sequencing a copy of each potential ancestor is calculated using the quality of reads of the nucleotides that differ between the two haplotypes: the score for each substitution address was calculated as the lowest quality over all reads sequenced in that address for the candidate haplotype. The final score was calculated from the product of all single substitution scores. The probability *P_ij_* that a candidate haplotype *i* was generated by sequencing errors in the reading of a potential ancestor *j* was determined using the complementary cumulative density (CCDF) of a binomial function, with the number of trials equal to the frequency of the potential ancestor, the number of successes equal to the frequency of the candidate haplotype and the success rate taken as the single event probability. This probability is calibrated according to the probability of ancestor *P_j_* so that 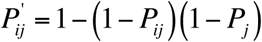. Finally, the P-value of haplotype *i* is calculated as the maximum probability over all potential ancestors: 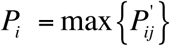.

### Calculation of probability of infection *P_infect_* from growth rates

To obtain the fitness landscape in **Figure 3**, we define the probability of infection of each haplotype as *P_inf_ = Θ(ρ-10^-3^)*, where *ρ* is the growth rate of the haplotype calculated as described below. Applying this threshold is motivated by the fact that the distribution of growth rates is bimodal – most haplotypes are either infective to some extent (*ρ*>=0.1) or not (*ρ*=0) with less than 1% of growth rates fall within the intermediate region (see **Figure 3-figure supplement 1A**). Consequently, the setting of the threshold is insensitive in the region (0,0.1).

### Calculation of growth rates from haplotype frequencies

We calculate growth rates by comparing between haplotype frequencies of consecutive generations. To avoid sequencing errors, we only consider frequencies with P-value≤10^-4^. Additionally, since our growth rates are relative to that of the wildtype *wt*, we only use data from the first 2 passages of evolution, where the *wt* is still the most abundant haplotype.

For each pair of consecutive generations that were sequenced, we calculate the growth rate *ρ*_i_ of all haplotypes that were detected in the samples using a simple linear model:

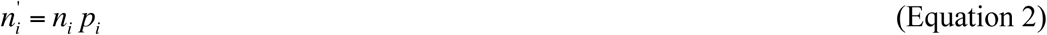

where *n*_i_ and *n*_i_' are the copy numbers of haplotype *i* in the first and second generation respectively. We ignore mutation rates since mutation rates of MNV (~10^-4^) are smaller than the sequencing errors (~10^-3^). Assuming unbiased sampling of original copies during sequencing, we substitute:

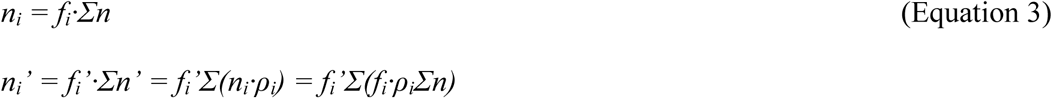

where *f_i_* and *f_i_*' are the frequency of haplotype *i* measured by sequencing the first and the second sample respectively. When we use these substitutions in the original equation (1) we get:

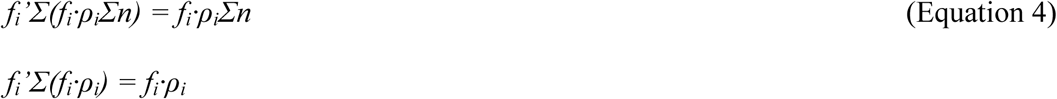

the resulting set of linear equations, one per haplotype, is a homogenous system:

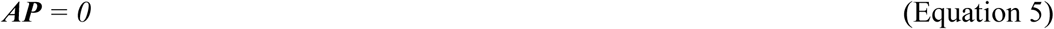

where the parameter matrix is *A_ij_* = *f_i_*'*f_j_*-*Δ_ij_* and the growth rates vector is ***P**_j_* = *ρ_j_*.

The non-trivial solution solves for all growth rates, assuming a growth rate of 1 for the wild type.

Since the same haplotype may exist in more than one pair of consecutive generations, multiple growth rates can be assigned to the same haplotype, in which case the average growth rate is calculated for this haplotype. The distribution of all growth rates is plotted in **Figure 3-figure supplement 1A**.

### Viral fitness function based on thermodynamics of protein folding and binding

We assume that the fitness *F* is proportional to fraction of MNV-1 P-domain that are folded and free from the antibody. If we further assume that the MNV-1 P-domain exists in three states— folded unbound, folded bound and unfolded (always unbound)— then we can express fitness *F* as

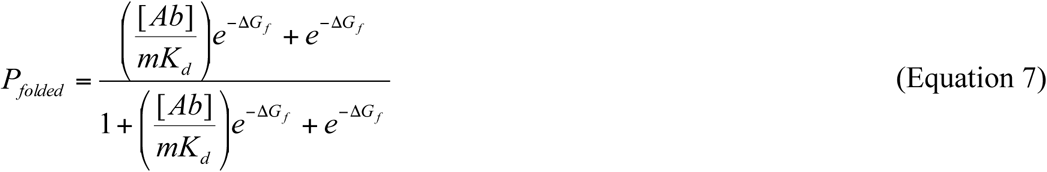

where Δ*G_fold_* and Δ*G_bind_* are the free energies of folding and binding, [*Ab*] is the antibody concentration, *C*_0_ is the standard reference concentration and *β*=1/*k_B_T*. The parameter *m* accounts for the added entropy due to the fact that any of the 180 P-domains on the viral capsid have a similar contribution to the neutralization of the capsid.

We note that this formulation takes into account the known experimental fact that binding induces folding. Specifically, the total fraction of folded proteins is the sum of proteins that are folded and bound to the *Ab* and proteins that are folded and free, and then normalized by total number of proteins in solution:

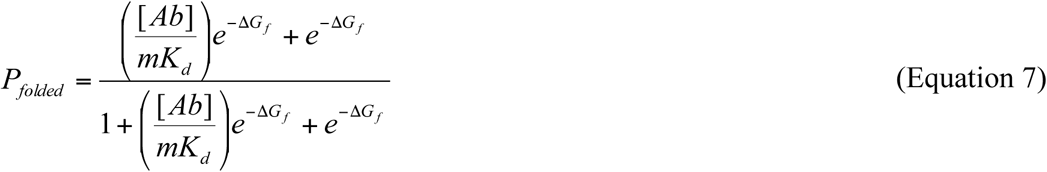

Indeed, *P_f_* → 1 when [*Ab*] → ∞ or *K_d_* → 0. That is, when there is a sufficient amount of antibody or when the binding to the Ab is very tight, all P-domains will eventually bind to the antibody and be folded.

### Fitness effects of mutations (selection coefficient) on the viral fitness landscape

Conceptually, evolution may be thought of as a hill-climbing process, whereby the rate of ascent is proportional to the effect of mutations on fitness. The fitness effect of a mutation is quantified by the selection coefficient *s*=(*F_mut_*-*F_wt_*)/*F_wt_*. The subscript *“mut”* and “*wt*” refers to mutant and wildtype, respectively. At the protein level, a non-synonymous mutation will change the folding and binding free energies as Δ*G_fold,mut_*=Δ*G_fold,wt_*+ΔΔ*G_fold_* and Δ*G_bind,mut_=ΔG_bind,wt_+ΔΔG_bind_*. For the fitness function (**Equation 1** and also **Equation 6**), we can derive a simple expression for the selection coefficient.

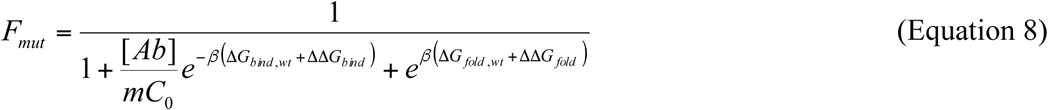

Consider the case when ΔΔ*G_fold_*<<1 and ΔΔ*G_bind_*<<1. Then, using the expansion *e^x^*≈1+*x*, when *x*<<1:

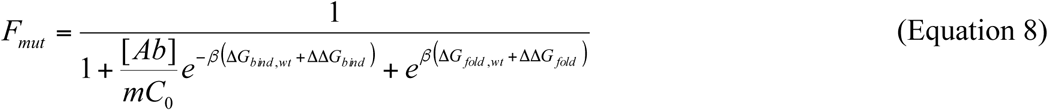

Thus, the selection coefficient is

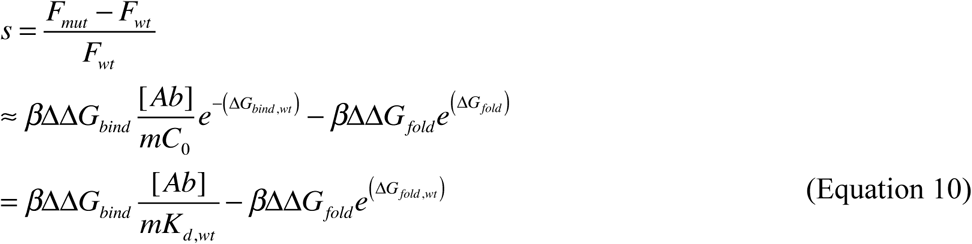

From **Equation 10**, the magnitude of the selection coefficient of a given mutation depends on the wildtype folding stability (Δ*G_fold,wt_*) and binding affinity (*K_d,wt_*); hence, there is strong epistasis on the fitness landscape. Moreover, because a single mutation can affect both folding and binding—mutations are pleiotropic in their effect to proteins—both terms in **Equation 10** contribute to the selection coefficient. This epistasis-pleiotropy link at the level of protein biophysics is the origin of the coupling between the selection for folding stability and binding.

The difference in sign for the two terms in **Equation 10** also denotes the opposite direction of selection for the two biophysical traits. Along folding stability, selection points towards more stable proteins (more negative values of A*G_fold_*); along binding affinity, selection points towards weaker binding (larger values of *Kd*) (**Figure 5A**). By contrast, randomly occurring mutations tend to destabilize proteins and protein-protein complexes, rendering both ΔΔ*G_fold_* and ΔΔ*G_bind_* more positive. Thus, along folding stability, mutational drift points towards less stable proteins, opposite to the selective force; however, along binding affinity, drift also points towards weaker binding (**Figure 5A**), parallel to the selective force. We present a formal treatment of the balance between selection and drift in the next section.

### Balance between selection and mutational drift on the fitness landscape: Theoretical analysis

We now formally combine the contribution of drift and selection and quantitatively determine the course MNV-1 evolution on the fitness landscape. We first limit ourselves to the simple monoclonal regime, where each generation is dominated by just one strain with a probability equal to its probability of fixation. To that end, we compare the probability of fixation of a non-synonymous mutation that changes folding stability and binding with the probability of fixation of a synonymous mutation, which we assume is neutral (s=0). We denote this ratio as *co.* The underlying motivation for this ratio is that when ω=1, mutations are neutral at the level of fitness, and mutational drift dominates. Specifically, ^51^

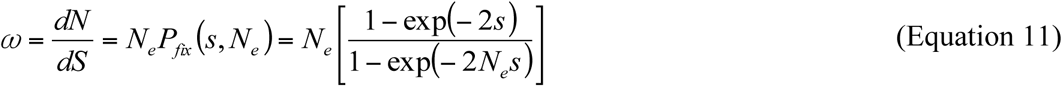

where *P_fix_*(*s*,*N_e_*) is the probability of fixation, *N_e_* is the population size and *s* is the selection coefficient (**Equation 10**). *s* is a function of the wildtype folding stability Δ*G_wt_* and binding affinity *K_0_* and the changes induced by the mutation to folding stability and binding ΔΔ*G_bind_* and ΔΔ*G_fold_*. Because the effect of random mutations on folding stability and binding, denoted by *p*(ΔΔ*G_fold_*) *an*d *p*(*ΔΔ_bind_*), are well-characterized ^38,39^ we can derive the rate o protein evolution averaged over all possible effect of random mutations on folding and binding, that is,

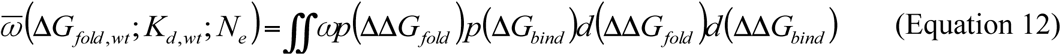

Eq. S10 defines the average stringency of Darwinian selection as a function of background, and hence the strength of epistatic interaction on the fitness landscape defined by protein folding and binding.

We show in **Figure 5A-B** (colormap) the value of the integral over the range of *K_d,wt_* and Δ*G_fold,wt_* values. Indeed, positive selection is strongest when the *wt* P-domain is unstable (low Δ*G*) or tightly bound to the antibody. At high population sizes, purifying selection and positive selection dominates in the regime of low folding stability and tight binding affinity (red colors in **Figure 5A**). The prediction is that if the population starts in the unstable regime, it would migrate away from the regime where selection is strong and towards the regime where it is more neutral.

### Simulations of evolutionary dynamics on the fitness landscape: Monoclonal regime

We complement our theoretical analysis above by performing explicit simulation of population dynamics. This evolutionary dynamics allows for the occurrence of random mutation and then selection. The algorithm is as follows:

#### Initialization

We start with a viral population of size *N.* Since the population is monoclonal, the entire population is uniquely determined by single values of folding stability and binding affinity. That is, the population is a single point defined by the ordered pair (Δ*G_fold_, K_d_*). For the simulation shown in **Figure 5A-B**, the population starts at (-2.5 kcal/mol, 46 nM).

#### Propagation

1. A non-synonymous mutation occurs in the MNV-p-domain at a rate of **µ*_r_* = 0.1 per replication ^52^. This mutation changes the folding and binding free energies as Δ*G_fold,mut_* = Δ*G_fold,wt_* + ΔΔ*G_fold_* and Δ*G_bind,mut_* = Δ*G_bind,wt_* + Δ*G_bind_*. The value ΔΔ*G_fold_* is drawn from a Gaussian distribution with mean 1 kcal/mol and s.d. .7 kcal/mol ^37,38^. The value ΔΔ*G_bind_* is drawn from a Gaussian distribution with mean 1 kcal/mol and s.d. 2 kcal/mol ^37,38^. Consequently, *K_d,mut_ = K_d,wt_e*^(βΔΔ*Gbind*)^.
2. Calculate the fitness of the mutant *F_mut_* using **Equation 1**.
3. Calculate the selection coecient *s* = (*F_mut_* – *F_wt_*)/*F_wt_*.
4. Determine the probability of fixation ^4^

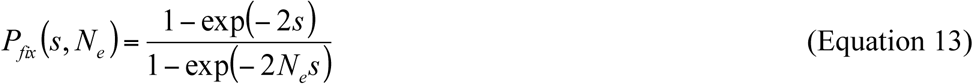
5. Draw a random number *r* between 0 and 1. If *r* <= *P_fix_*, the mutation is fixed and the folding stabilities and binding affinities are updated; otherwise, keep the wildtype values.
6. Repeat steps 1-5 for a specified number of fixations (In **Figure 5A-B**, we ran simulations until fixation of 4 amino acid substitutions).

### Simulations of evolutionary dynamics on the fitness landscape: Polyclonal regime

The analyses in the previous section are performed in the monoclonal regime, where a closed formula of the stringency of selection is tractable. However, a true viral quasi-species is necessarily polyclonal because of the high mutation rate, thus, we also perform forward polyclonal regime simulation, for which the closed form formula for the stringency of selection is no longer accurate. Results are shown in **Figure 5-figure supplement 1**.

#### Initialization

We start with a viral population of size *N.* Each of the *N* virion in the population is characterized by its folding stability and binding affinity. Thus, the entire population constitutes a distribution of Δ*G_fold_* and *K_d_* values and migrates as a “cloud” on the fitness landscape. For the simulation shown in Fig. S4, we start with a population of *N* randomly chosen variants from the sequenced MNV-1 viral stock.

#### Propagation

1. Each of the *N* viruses has a chance to infect and replicate with a probability given by the fitness function **Equation 1**.
2. Each replicating virus mutates at a rate of **µ*_r_* = 0.1 per p-domain per replication ^52^. If a mutation occurs, the folding stability and binding affinity are updated similar to Step #1 in the monoclonal regime. The number of progenies is calibrated to *N* to maintain a constant population size.
3. If no viruses are chosen for replication, the sample is extinct.
4. Repeat from steps 1-3 for desired number of generations. To compare the number of generations to the number of fixations in the monoclonal simulations, we assume that each fixation requires 4 generations.

## Acknowledgements

The authors would like to acknowledge C. Wilke and J. Plotkin for very helpful comments. This work was supported by Defense Advanced Research Projects Agency (DARPA) contract # HR0011-11-C-0093, by the National Science Foundation (DMR-1310266 to DW and MCB-1243837 to ES), NIH GM068670 (ES), by the Harvard Materials Research Science and Engineering Center (DMR-1420570) and by the National Natural Science Foundation of China (HZ, 81372496).

## Footnotes

**Competing Interests** The authors declare none.

---

*The following supplements are available for Figure 1:*

**Figure 1-figure supplement 1.**
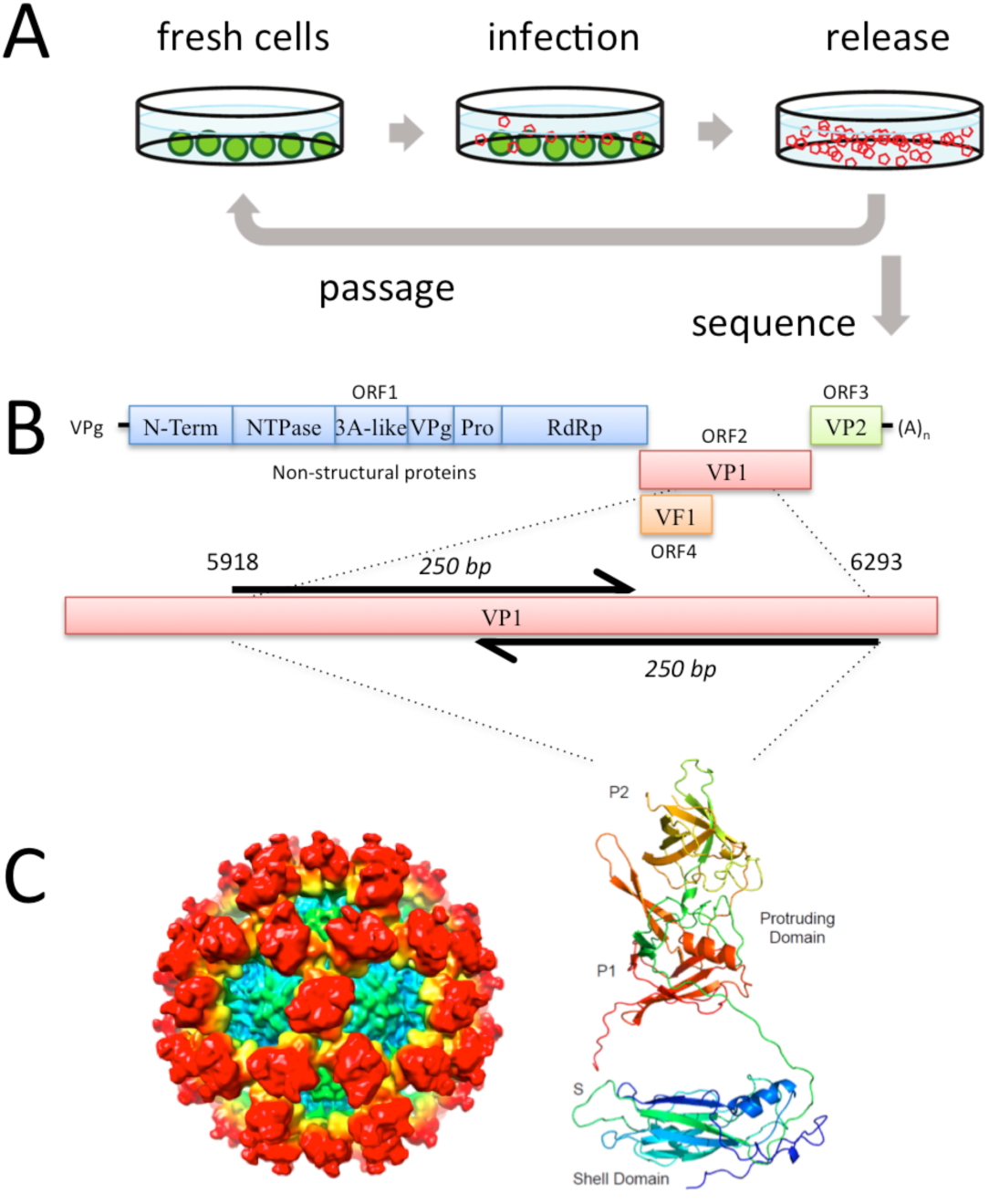
Overview of the experimental set-up. **A)** RAW 264.7 cells exposed to MNV-1 virus are infected and viral progenies are released into the supernatant. To passage viruses, a small volume fraction of the supernatant is transferred to a new sample of fresh cells. The viral genomes in the remaining supernatant are prepared for sequencing. **B)** The 7.5kb long MNV-1 genome includes 4 main reading frames with the capsid protein gene (VP1) residing in the second frame. For our purposes, we sequence a 376bp long fragment in VP1 that encodes part of the protruding domain (P domain) of the capsid protein. **C)** *Left:* Cryo-EM structure of MNV capsid, adapted from ^21^. *Right:* Model of MNV made by fitting the P-domain of MNV-1 ^24^ and the shell domain from the Norwalk virus structure^25^ into the cryo-EM density of MNV-1 ^22^.

**Figure 1-figure supplement 2.**
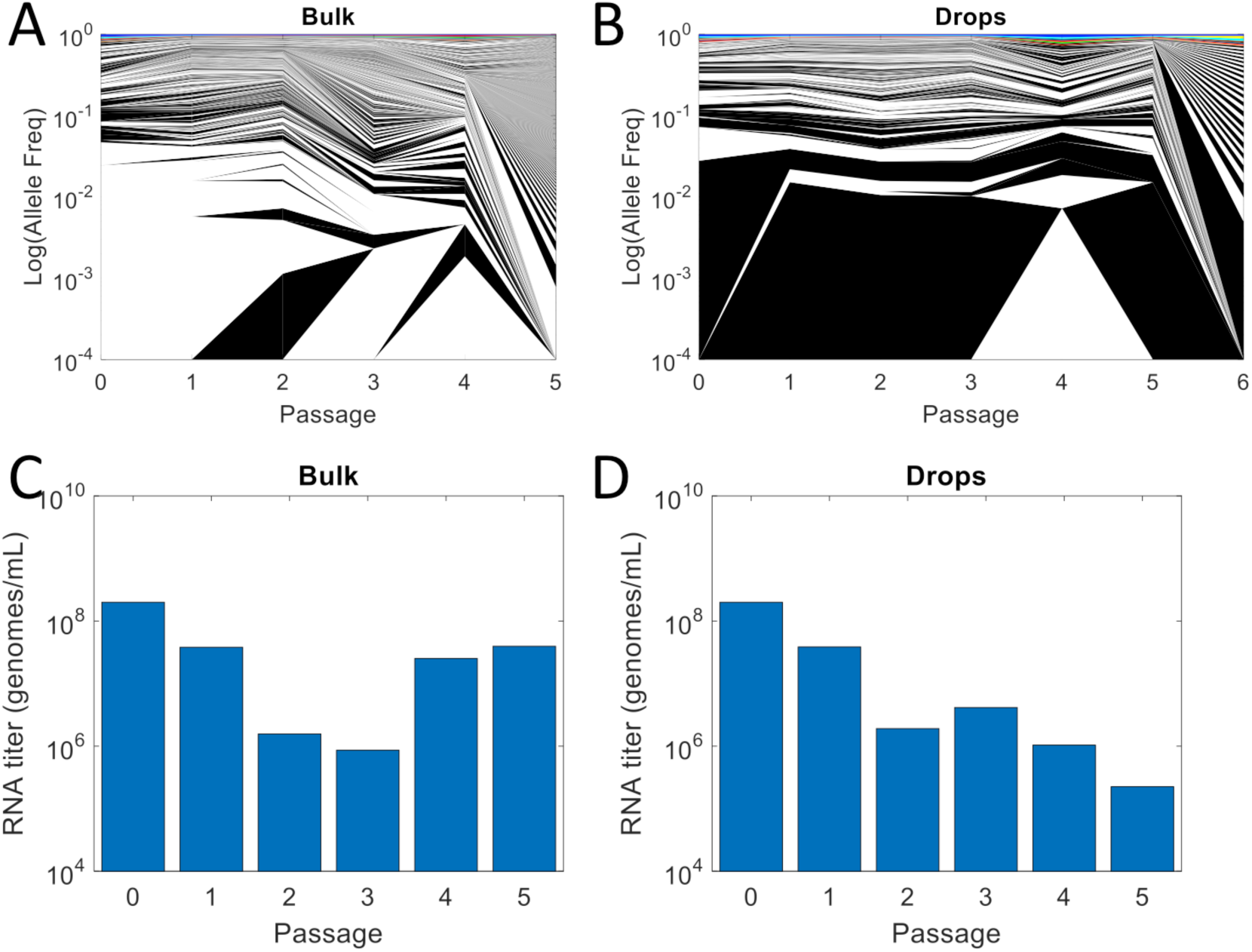
Allele frequency and viral RNA concentration from experimental viral evolution in bulk and drops. **A)** The allele frequencies of 1,364 distinct P-domain haplotypes of the evolution in bulk (**Figure 1A**) are plotted per passage, with the y-axis plotted in log-scale to show the polyclonal structure of the viral quasi-species. **B)** The allele frequencies of 620 distinct P-domain haplotypes of the evolution in drops (**Figure 1B**) are plotted per passage, with the y-axis plotted in log-scale to show the polyclonal structure of the viral quasi-species. We note that in (A) and (B), similar to primary **Figure 1**, dominant haplotypes are individually color coded but do not appear due to log transformation of Allele frequency. **C, D)** The concentration of viral RNA in the experimental samples is plotted per passage for both the evolution in bulk (C) and that in drops (D).

*The following supplement is available for Figure 2:*

**Figure 1-figure supplement 3.**
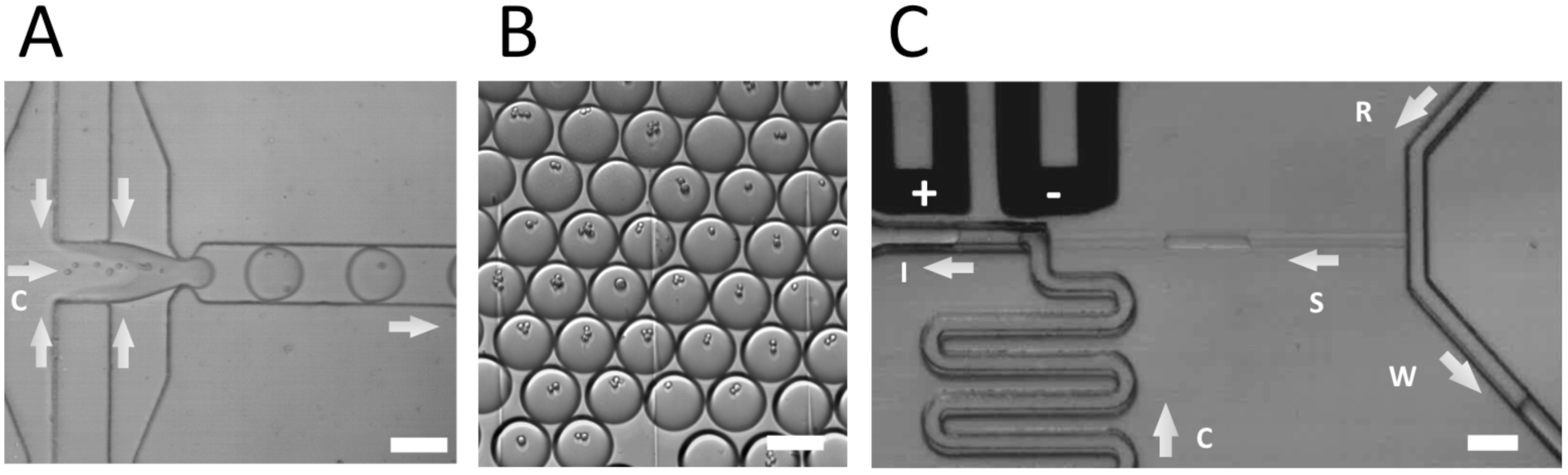
“Evolution Chip”: a system for evolving virus in very small populations. *Left:* Virus and host cells co-encapsulation. *Middle:* Drop incubation. *Right:* Drop passaging. All scale bars are 100 *µ*m.

*The following figure supplements are available for Figure 3:*

**Figure 3-figure supplement 1.**
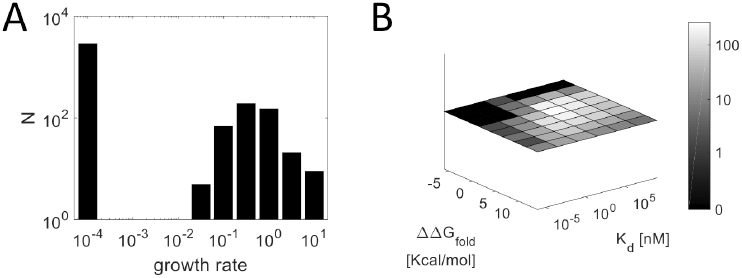
Distribution of viral fitness estimated from deep sequencing. **A)** Distribution of 3,770 growth rates calculated from the first 3 passages of all evolution experiments. **B)** The number of haplotypes that were binned in each 2D bin of **Figure 3D** is color coded with a logarithmic gray scale representation.

**Figure 3-figure supplement 2.**
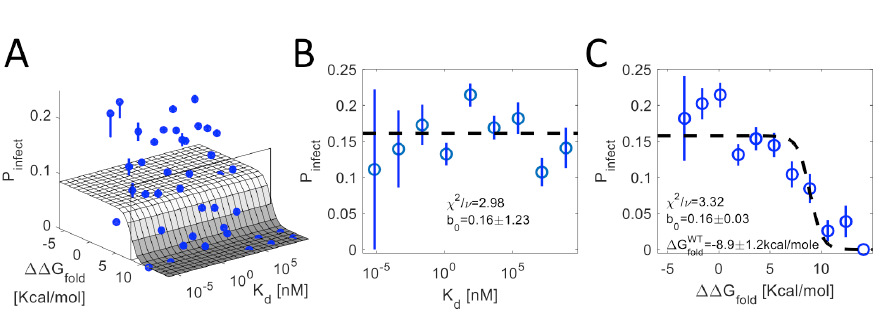
Fitness landscape of MNV-1 in the absence of a neutralizing antibody. **A)** A 3D plot of the probability of infection (z-axis) averaged over ~2,000 distinct haplotypes binned according to their dissociation constant *K_d_* and folding stability ΔΔ*G_fold_* (blue error bars) overlaid with the theoretical fit according to Equation 1 (gray surface). Cross sections (black boxes) border the regions used for the projections in B and C. **B)** The probability of infection for all haplotypes with ΔΔ*G_fold_* < 8*kcal*/*mol* (cross section parallel to *K_d_* axis in A) projected on the *K_d_*-z plane, binned according to their *K_d_* (blue error bars) and overlaid with the theoretical fit according to Equation 1 (dashed line). **(C)** The probability of infection for all haplotypes is projected on the ΔΔ*G_fold_*-z plane, binned according to their ΔΔ*G_fold_* (blue error bars) and overlaid with the theoretical fit according to Equation 1 (dashed line). Probability of infection is determined from deep sequencing lysates of *in vitro* experiments in the absence of neutralizing antibody *K*_d_ and ΔΔ*G_fold_* are estimated from mapping the haplotypes' mutations to the 3D structure of the capsid P-domain in complex with the neutralizing antibody.

*The following figure supplement is available for Figure 4:*

**Figure 4-figure supplement 1.**
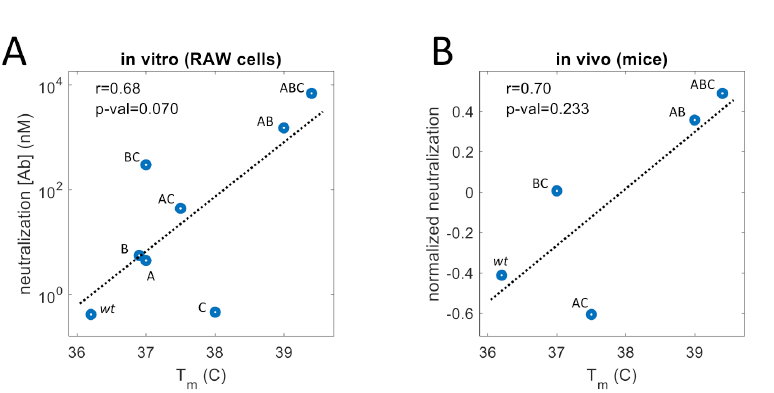
MNV-1 neutralization versus folding stability of the P-domain. **(A)** MNV-1 neutralization in RAW cells correlates with *T_m_*. Using reverse engineering, we introduce the dominant SNPs (**Figure 3A**) on the background of *wt* MNV-1 and measure their effective neutralization (see *Methods*). **(B)** MNV-1 neutralization in mice weakly correlates with *T_m_* (see *Methods*).

*The following figure supplement is available for Figure 5:*

**Figure 5-figure supplement 1.**
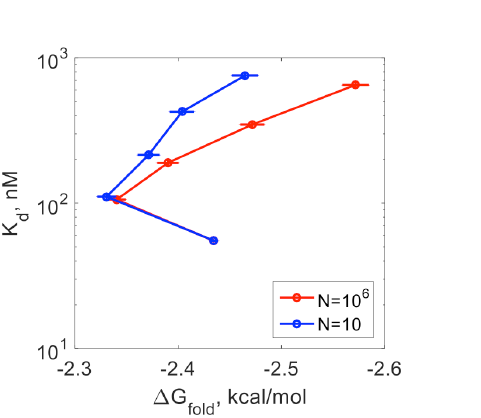
Simulations of evolutionary dynamics on the fitness landscape with a polyclonal viral population. The average folding stability and *K_d_* is shown for 5 simulated passages of populations of 10 and 10^6^ viruses. The simulation was repeated once for the large population and 1,000 times for the small population and the average of all repeats are shown (SSE error bars are not visible at < 0.1%). For this simulation we used the initial conditions: Δ*G_fold_* =-2.5kcal/mol, *K_d_* =46 nM, [*Ab*]=8 nM and **µ*_r_*=0.1.

**Table S1.** Complete list of MNV-1 variants from deep sequencing

**Movie S1.** Encapsulation of virus and cells in drops

**Movie S2.** “Evolution Chip” for serial passaging in drops

**Table S2.** Analysis for head to head competitions

